# Low-threshold, high-resolution, chronically stable intracortical microstimulation by ultraflexible electrodes

**DOI:** 10.1101/2023.02.20.529295

**Authors:** Roy Lycke, Robin Kim, Pavlo Zolotavin, Jon Montes, Yingchu Sun, Aron Koszeghy, Esra Altun, Brian Noble, Rongkang Yin, Fei He, Nelson Totah, Chong Xie, Lan Luan

## Abstract

Intracortical microstimulation (ICMS) enables applications ranging from neuroprosthetics to causal circuit manipulations. However, the resolution, efficacy, and chronic stability of neuromodulation is often compromised by the adverse tissue responses to the indwelling electrodes. Here we engineer ultraflexible stim-Nanoelectronic Threads (StimNETs) and demonstrate low activation threshold, high resolution, and chronically stable ICMS in awake, behaving mouse models. *In vivo* two-photon imaging reveals that StimNETs remain seamlessly integrated with the nervous tissue throughout chronic stimulation periods and elicit stable, focal neuronal activation at low currents of 2 μA. Importantly, StimNETs evoke longitudinally stable behavioral responses for over eight months at markedly low charge injection of 0.25 nC/phase. Quantified histological analysis show that chronic ICMS by StimNETs induce no neuronal degeneration or glial scarring. These results suggest that tissue-integrated electrodes provide a path for robust, long-lasting, spatially-selective neuromodulation at low currents which lessen risks of tissue damage or exacerbation of off-target side-effects.

## Introduction

Built upon the success of electrical stimulation from macroelectrodes to induce coarsely focused cortical activation(Fritsch and Hitzig, 1870; Penfield and Boldrey, 1937) intracortical microstimulation (ICMS) using implanted microelectrodes modulates neural activity and elicits behavioral responses at finer spatial resolutions (Asanuma and Sakata, 1967; Ekstrom et al., 2008; Otto et al., 2005; Stoney et al., 1968; Tabot et al., 2013; Venkatraman and Carmena, 2011). In ICMS, intracortically implanted microcontacts inject electrical charges into the surrounding tissue, create flow of ionic current, depolarize the membranes of excitable cells, and change neural activity locally. This capability enables diverse applications such as establishing causal links between neural activity and behavior(Asanuma and Sakata, 1967; Gallistel et al., 1981; Tehovnik, 1996), modulating attention and learning(Voigt et al., 2018), and producing perception and sensations (Armenta Salas et al., 2018; Chen et al., 2020). Across all these diverse applications, the overall technological goal of ICMS is to produce targeted, high-resolution neuronal modulation capable of eliciting stable perception or sensation over an extended period(Pancrazio et al., 2017; Urdaneta et al., 2017). Because ICMS often induces neuronal activation from the passage of axons(Histed et al., 2009), realizing this goal requires i) subcellular proximity and stability at the tissue-electrode interface, and ii) the stimulation electrode to reliably produce identical, highly localized charge injections.

Current ICMS electrodes are significantly more rigid than the host brain tissue, resulting in instability of the tissue-electrode interface and substantial “spatiotemporal blur” in the neuronal response(Butovas and Schwarz, 2003). Over chronic implantation durations, the tissue-electrode interface deteriorates(Grill et al., 2009; Polikov et al., 2005). Neuronal degeneration and formation of glial scarring around the probe alters the electric fields induced by the stimulus, which could change the resultant neural(Davis et al., 2012) and behavioral(Koivuniemi et al., 2011) responses. Critically, related to the interface degradation, large and increasing amplitudes of stimulation are often needed to maintain behavioral responses in chronic applications(Davis et al., 2012; Koivuniemi et al., 2011), which expedites the deterioration of stimulating electrodes(Cogan et al., 2004) and increases the risk of tissue damage(McCreery et al., 2010; McCreery et al., 2002). Furthermore, significant longitudinal variations have been reported, but it is not clear whether they are due to changes in adverse tissue responses, degradation of electrodes, or intrinsic changes in the excitability or functional response of the neural tissue.

Theoretically, promoting device-tissue integration will enhance electrode-neuron proximity and therefore improve the resolution and stability of ICMS. Based on fundamental biophysical principles, implants with no glial scar encapsulation will minimize the separation between the electrode and targeted neurons, which will reduce the activation threshold, decrease the number of activated neurons at threshold, improve focality, eliminate time-varying foreign-body tissue responses, and result in high-resolution, chronically stable neuromodulation (**Figure 1A**). We performed proof-of-principle finite-element-model simulations to compare the activation threshold of neurons at various glial scar thickness (0, 20, and 40 μm). The absence of glial scarring reduces the current required to activate the same number of neurons and lowers the tissue volume of activation (**Figure 1B**). Intuitively, this effect scales with the thickness of the glial scar: the thinner the scar, the smaller the current required to stimulate neurons and the lesser the volume of tissue activation near the threshold (**Figure 1B**). Furthermore, reducing the glial scar improves the spatial selectivity of stimulation. When stimulating two nearby contacts, reducing the thickness of the scar layer reduces the spatial overlap of stimulation. Closely spaced stimulation sites with no scar encapsulation provide the most spatially distinct tissue activation compared with those with scars at the same tissue activation volume (**Figure 1C**).

**Figure 1:**
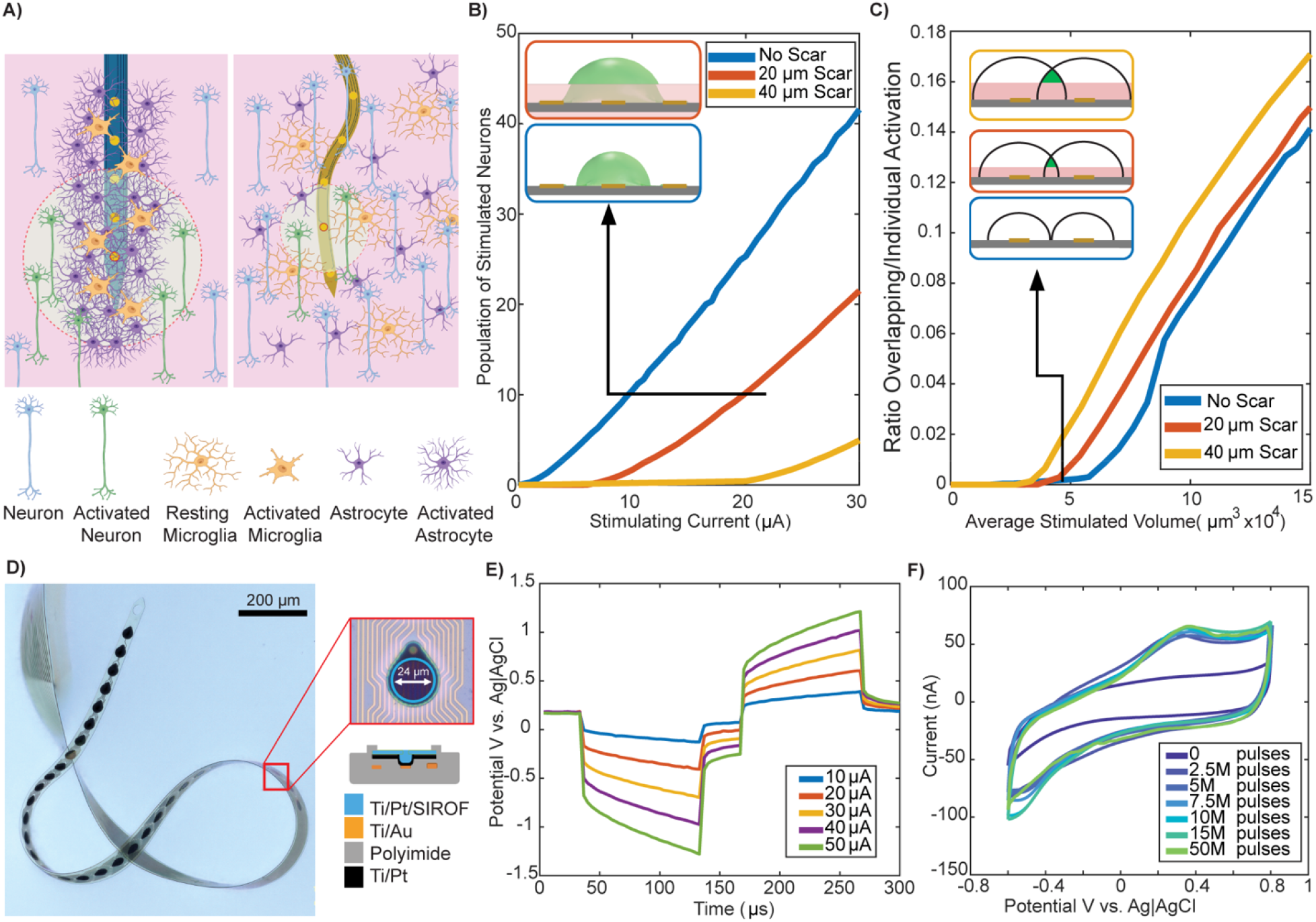
Engineering tissue-integrated flexible electrodes to enhance microstimulation efficacy. **(A)** Diagram of chronic immune response to rigid and flexible implants showing that, in theory, no glial scarring reduces the distance between the stimulation site and neurons so that a lower current can elicit more focal neural activation. **(B)** Simulation showing that the current needed to activate the same number of neurons reduces with glial scar thickness. Inset: volume of the neuronal tissue (shade green) when activating 10 neurons in the situation of no scar (bottom) and 20 μm thick scar (top). Gray: polyimide; Golden yellow: stimulation contacts. Shaded pink: glial scar. **(C)** Simulation showing that overlapping volume increases with scar thickness when stimulating two nearby sites. Inset: spatial profile of the stimulated tissue volume when one stimulation site activated the same volume in the scar thickness of 0, 20, and 40 μm. Black: boundary of activated tissue by each stimulation site. Shaded green: overlapping region. **(D)** Photo of a meandering StimNET in water showing ultra-flexibility. Inset: zoom in photo of a stimulation site and its cross-sectional structure. SIROF: Sputter IrO_x_ film. **(E)** Representative in-vitro voltage transients at various current amplitudes. **(F)** In-vitro cyclic voltammograms at 100 mV/s showing stable charge storage capacity of Stim-NETs after 50 million pulses at 30 μA.

The biophysical principals and results from numerical simulations motivated us to maximize the efficacy of ICMS by optimizing the tissue-electrode interface. We engineered the device flexibility and developed the ultraflexible stim-Nanoelectronic Threads (StimNETs) to meet the requirements of robust charge injection and subcellular stability at the tissue-electrode interface simultaneously. We employed a suite of optical, electrical, behavioral, and histological methods in mouse models to evaluate the efficacy, resolution, stability, and tissue compatibility of neuromodulation. We verified that these tissue-integrated electrodes produce spatially confined neuronal activation and elicit longitudinally stable behavioral detection at substantially reduced stimulation currents with no neuronal degradation or glial scarring. These results underscore the importance of tight tissue-electrode integration in the efficacy of stimulation and provide a path for long-lasting, high-resolution neuromodulation at low currents that minimize risks of tissue deterioration or exacerbation of off-target side-effects.

## Results

### Engineering ultraflexible StimNETs for robust charge injection

We chose to drastically reduce the substrate thickness of intracortical electrodes to minimize the bending stiffness and provide tight tissue-electrode integration. Our previous study demonstrated that Nanoelectronic Threads (NETs) at a total thickness of 1 μm forms an intimate tissue-electrode interface during chronic implantation, featuring an intact brain-blood barrier, tissue-electrode stability at subcellular scale for over a few weeks, and an absence of neuronal degradation and glial scarring near the electrodes(Luan et al., 2017; Wei et al., 2018). However, the ultrathin insulation layer (0.5 μm) and multi-layer device architecture impose significant challenges for lasting stimulation without structural and functional breakdown. Through iterative testing and device optimization we have realized ultraflexible StimNET for robust stimulation at the similar form factors and ultraflexibility as the recording NETs (**Figure 1D**). We focused on the following modifications. First, to reduce the risk of crosstalk between nearby trace lines, we switched the substrate material from SU8 to polyimide (PI), which is a stronger dielectric with larger tensile strength, and adapted multi-layer planar microfabrication on PI (STAR Methods). Second, to improve the charge storage capacity and charge injection capacity of NET microcontacts, we sputtered IrO_x_ on Au contacts at the wafer scale during microfabrication(Slavcheva et al., 2004). Third, as a precaution to cover potential cracks on IrO_x_ vertical walls, we offset vias and contacts. Lastly and importantly, to alleviate the risk of delamination, we microfabricated a cap ring (thickness of 0.3 μm) using PI surrounding each contact on top of IrO_x_ as an additional mechanical reinforcement (**Figure 1D**). The optimized StimNET has a shank thickness of 1 μm and an additional 0.3 μm thickness at the cap ring, interconnect linewidth of 1.3 μm, and 32 individually addressed microcontacts per shank for both recording and stimulation at a diameter of 24 μm.

To verify the charge injection and storage capacity of StimNETs, we performed voltage transient measurements and cyclic voltammetry (CV) in vitro (STAR Methods). Individual microcontacts in Stim-NET output currents up to 50 μA while maintaining the maximum cathodically and anodally driven electrochemical potential excursion within the water window of [−0.6 – 0.8 V] (**Figure 1D**). The charge injection capacity was on par with rigid electrodes with sputtered IrO_x_(Cogan, 2008). **Figure 1F** shows a representative pulsing test in which we stimulated a contact site at 500 Hz, 30 μA biphasic charge-balanced pulses for 50 million pulses and acquired CV at a scan rate of 100 mV/s periodically during pulsing. Except for an initial increase in charge storage capacity, which is well documented as increased porosity and accessibility of Ir^3+^/Ir^4+^ redox sites with continual oxidation(Maeng et al., 2020), there was little change in CV between 2.5 million to 50 million stimulation pulses. These results demonstrated the high charge injection and storage capacity of StimNETs and supported their durability and robustness during stimulation.

### High-throughput quantification of ICMS-evoked neuronal activation in awake animals

To evaluate ICMS efficacy over a chronic period at single-cell resolution, we co-implanted a cranial window and StimNET in the somatosensory cortex of Thy1-GCamp6s mice and performed two-photon (2P) Ca^2+^ imaging during ICMS (**Figure 2A,** Methods). Different from most prior studies of the spatial activation pattern of ICMS which used anesthetized animals(Histed et al., 2009; Michelson et al., 2019), we performed 2P z-stack imaging of neuronal activations in awake animals to remove confounding effects of anesthesia (STAR Methods). **Figure 2B** shows a set of representative Ca^2+^ images across the cortical depth acquired during stimulation where cells were fluorescent either due to spontaneous or ICMS-evoked activation. To quantify the ICMS-evoked neuronal activation from the background of spontaneous activity, we used a trial structure where the randomized ICMS trials alternated with baseline trials (spontaneous activity, no stimulation) for differential measurements and analysis between each paired stimulated and baseline trials (STAR Methods). Using galvo resonant scanners and customized automatic data acquisition pipelines, we were able to acquire 300 – 400 z-stacks of 1 mm × 1 mm × 0.4 mm in a typical 3-hour session that contained blocks of randomized stimulation channels and currents, and 7 – 9 replicas of these blocks. Representative trials of four current levels and three stimulation channels showed clear modulation of Ca^2+^ fluorescence (**Figure 2C**, trial sequences were recognized and grouped by currents and stimulation channels for presentation clarity).

**Figure 2:**
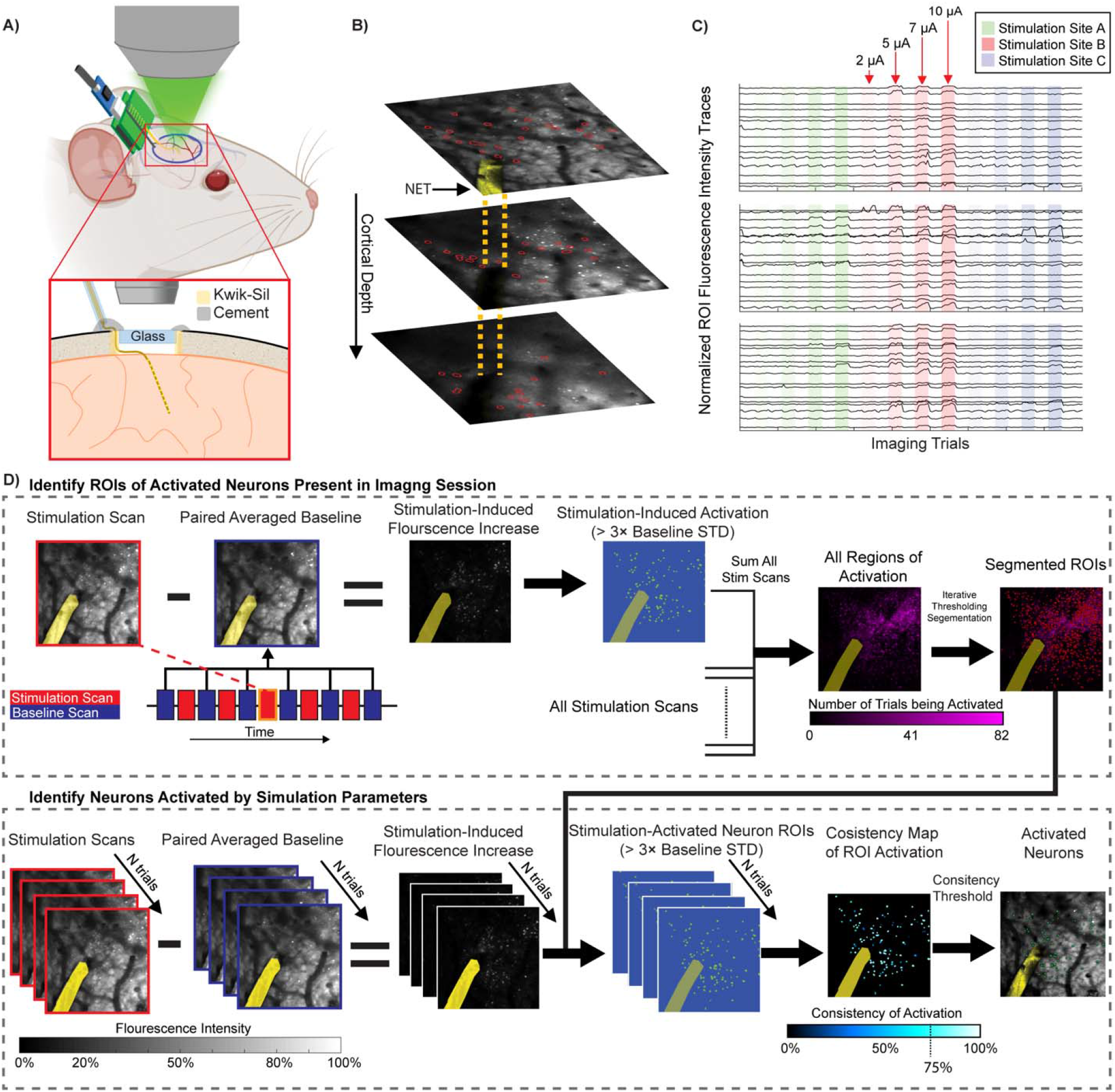
High-throughput quantification of ICMS-evoked neuronal activation in awake animals. **(A)** Diagram illustrating the surgical preparation and experimental setup for synchronous two-photon Ca^2+^ neural imaging and ICMS in awake mice for longitudinal studies. **(B)** Representative Ca^2+^ images across the cortical depth during ICMS. Yellow ribbon and dashed lines: image and sketch of the StimNET. Red circles: representative neurons shown in (C). **(C)** Sample fluorescence intensity traces from neurons in b over multiple stimulation and baseline trials. A distinct subset of cells was activated by each stimulation site. **(D)** Image processing workflow to identify and localize neurons activated by ICMS in awake animals. Regions of interest (ROIs) for active neurons were identified by differential measurements of stimulated and non-stimulated trials, thresholding against baseline fluorescence variance, and segmented for the entire imaging session. The segmentation results were then fed to the activation map of each stimulation parameter, which, after checking for consistency of activation > 75% across repetitive trials, isolated the activated neurons by this specific stimulation parameter. Yellow ribbon: StimNET. Red dots: neuron ROIs. Green dots: evoked neurons.

We then developed and used computationally efficient matrix manipulations to calculate the difference between paired stimulation and baseline, identified the voxels that were activated by any current and stimulation sites, quantified the probability of activation across N repeats, and imposed a probability threshold (here we set as 75%) across all trials to identify areas that were consistently activated only upon stimulation (**Figure 2D**). The most computational demanding step, segmentation to identify individual neurons evoked by ICMS was only performed once aggregating all trials in the same imaging section together. This pipeline greatly improves the throughput of imaging processing, making it feasible to identify individual neuron and their location in three dimensions (3D) under numerous stimulation parameters (STAR Methods). Additionally, because the activation regions are identified all together for all trials that use different stimulation sites and at various ICMS currents before segmentation, the segmented neurons are less prone to small drifts, which facilitate the comparison of neuronal activation pattern across stimulation sites and currents (STAR Methods).

### StimNET elicits spatially localized neuronal activation at low currents

To map the 3D spatial distribution of neuronal activation, we stimulated individual sites of StimNET in layer 2/3 in the somatosensory cortex, performed concurrent 2P z-stack imaging during ICMS up to 500 μm deep into the tissue, and identified the evoked neurons and their locations. We detected activation of a small number of cells near the electrodes at a low ICMS current of 2 μA (**Figure 3A**). Intuitively, the number of ICMS-evoked neurons increased with ICMS currents (**Figure 3B,** Kruskal-Wallis test with Dunn’s post hoc correction, degree of freedom (df)=3, 2 μA vs. 5 μA, p = 3.7e-9; 2 μA vs. 7 μA, p = 3.7e-9; 2 μA vs. 10 μA, p = 3.7e-9). The ability to identify each activated neuron in 3D allowed us to determine the ratio of neurons consistently activated at both low and higher currents when stimulated by the same site. We obtained a ratio of more than 85% for consistent activation at all currents (**Figure 3B**), which confirms the short-term stability of the tissue-electrodes interface owing to the mechanical compliance of StimNETs. Furthermore, we quantified the volumetric neural activation density (**Supplementary Figure S1**) as a function of currents and projected the 3D activation density into 2D for visualization (**Figure 3C,** averaging 5 animals, 11 imaging sessions, and 21 stimulation sites). The neuronal activation was highly localized near the stimulation site at 2 μA, a low current that was rarely studied previously. Once the current increased, the activation pattern became spatially distributed as previously reported at similar magnitudes of currents. Consistently, the spatial extent of ICMS-evoked neural activation, defined as the largest distance from any evoked neuron to the stimulation site, increased significantly from 2 μA to 5 μA (**Figure 3D**, Kruskal-Wallis test with Dunn’s post hoc correction, df = 3, p = 0.035), but remained relatively unchanged with further increase of ICMS currents (7 μA vs. 10 μA, p = 0.90). These experimental results supported our simulation (**Figure 1B**) and confirmed that the ultraflexible StimNET could elicit focal activation at very low currents.

**Figure 3:**
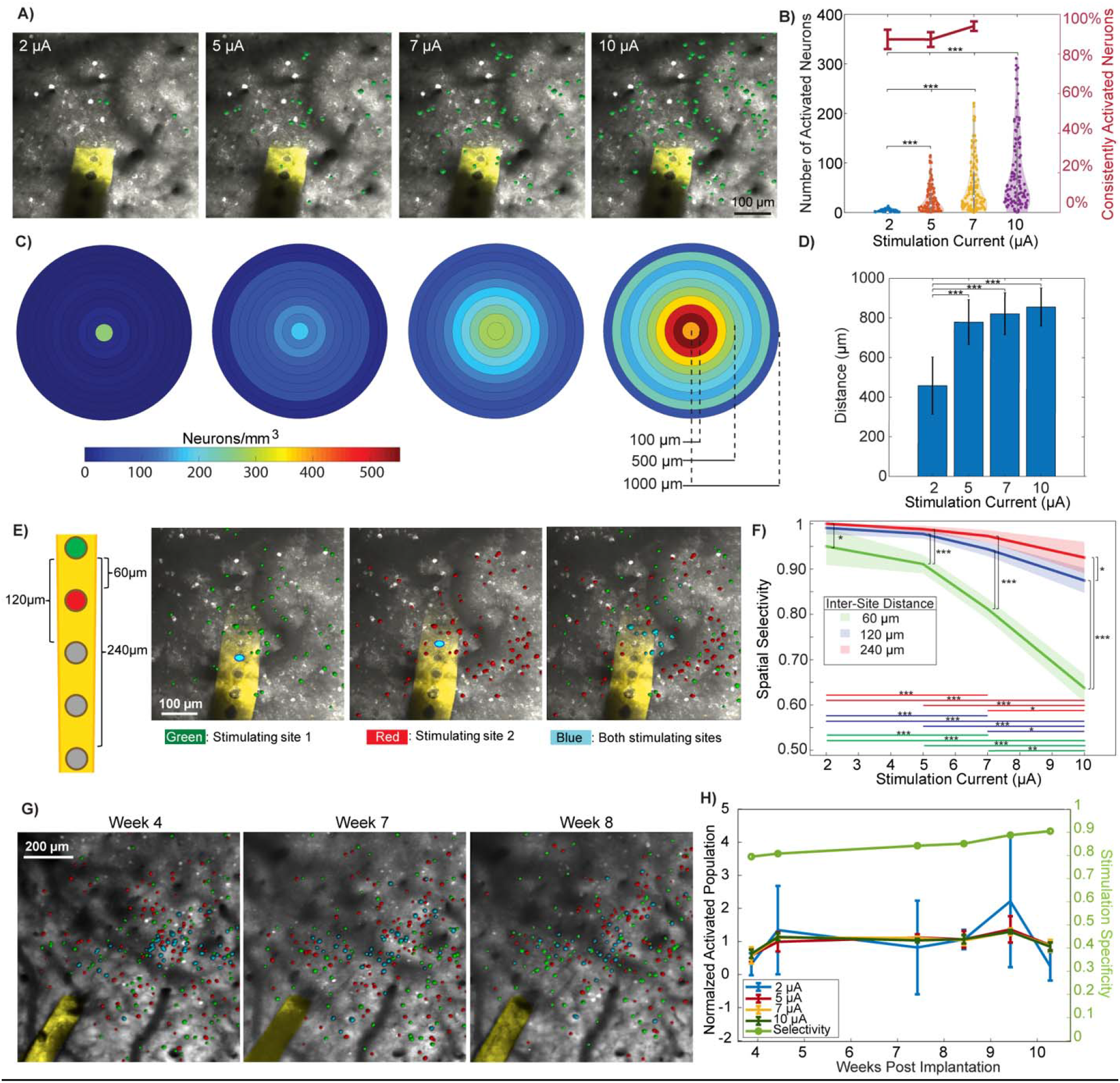
StimNET elicits spatially localized neuronal activation at low currents. **(A)** Representative 2P images showing neuronal activation increased with stimulation currents. Images are maximum intensity projections (MIP) of z-stack from 0 – 400 μm. Green: ICMS-evoked neurons; Yellow: StimNET. (**B**) Left axis: Violin plot of total neural population activation. Embedded whisker plots denote the 25^th^, 50^th^, and 75^th^ percentiles. Right axis: percentage of neurons being activated by the next higher current. Error bars denote 95% confidence intervals. **(C)** Ring plots showing the averaged cell activation density as a function of distance in 3D from the stimulation sites. (**D**) Bar plot of the maximum spatial spread of neural activation showing a significant increase from 2 μA to higher currents. **(E)** Representative 2P MIP of the same imaging volume showing adjacent stimulating sites at 5 μA activated distinctive populations. Green: neurons activated by site 1; Red: neurons activated by site 2; blue: neurons co-activated by site 1 and 2. Sketch on the left shows the site separation. **(F)** Spatial selectivity as a function of stimulation currents for three inter-site separations. (**G**) Representative 2P MIP in the same animal showing consistent and spatially selective activation of neurons over time. Same color code as in (E). Center-to-center distance of two neighboring sites: 60 μm; stimulation current: 7 μA. **(H)** Normalized neural activation shows stable population recruitment over time (left axis). Error bars denote 95% confidence interval. Right axis displays the spatial selectivity over time at 5 μA. Sample numbers: n=5 animals, 21 stimulation channels, and 11 imaging sessions for (B – D); n=6 animals, 79 stimulation channels, and 20 imaging sessions for (F). Statistical significance: * p<0.05; ** p< 0.01; *** p<0.001.

### Neuronal activation is spatially selective and numbers of evoked neurons are longitudinally stable

Most applications of ICMS will gain from the ability to activate discrete groups of neurons by neighboring contacts over an extended period. Therefore, we evaluated the spatial selectivity of StimNET, the ratio of the distinctive population activated by each contact over the total activation population, through longitudinal 2P Ca^2+^ imaging. (Method). Spatial selectivity would be 1 if two nearby contacts result in entirely different, non-overlapping activation of neurons, while selectivity would be 0 if they activate the exact same population. **Figure 3E** shows a representative neuronal activation pattern when stimulating two spatially adjacent contacts (center-to-center distance 60 μm) at 5 μA. Each contact elicited distinctive neuronal activation, and only a small number of cells were co-activated by both contacts. To quantify how spatial selectivity changes with ICMS current and separation of stimulating contacts, we stimulated while simultaneously imaging volumetrically at four levels of currents and three contact distances. At a low stimulation current of 2 μA, the spatial selectivity was >95%, even for contacts that were separated by merely 60 μm center to center (36 μm edge to edge). As expected, increasing stimulating currents significantly decreases selectivity for all the contact spacing tested (**Figure 3F**: Kruskal-Wallis test, χ^2^ = 136.24, p = 2.4e-29, df = 3). Particularly, the lowest current has the highest spatial selectivity (Kruskal-Wallis test with Dunn’s post hoc correction, 2 μA vs. 5 μA, p = 0.0002; 2 μA vs. 7 μA, p = 3.7e-9; 2 μA vs. 10 μA, p = 3.7e-9). For any given stimulation current, increasing the contact separation improves selectivity (Kruskal-Wallis tests with Dunn’s post hoc correction, df = 2; 60 μm vs. 240 μm at 2 μA, p = 0.02; and at 10 μA, p = 9.8e-12). Notably, a 120 μm center to center separation was sufficient to achieve > 90% spatial selectivity for all currents tested, which validates the application of StimNET for highly selective neuromodulation.

We repeated Ca^2+^ imaging longitudinally during stimulation until the cranial window got cloudy, mapped the neuronal activation spatially, and quantified the number of neurons being activated and the spatial selectivity when stimulating two neighboring contacts as a function of time after StimNET implantation. The spatial patterns of neural activation evoked by the same stimulation sites and currents were similar across time, while the actual neurons activated from week to week by each contact had mild changes (**Figure 3H**). The number of neurons activated at the same current and the spatial selectivity at the same stimulation parameters remained stable through the entire experimental duration (**Figure 3G**). These results suggest that over chronic periods the same currents from StimNET activated similar numbers of neurons and maintain the same high level of spatial selectivity.

### StimNET elicits robust, chronically stable behavioral detection at low currents

To evaluate the behavioral detectability of low amplitude ICMS via chronically implanted StimNETs, we developed and used a go/no-go task, for which water-deprived, head-fixed mice were trained to turn a wheel past an angular displacement threshold in response to ICMS to obtain water rewards (STAR Methods, **Figure 4A**). A random inter-trial interval (2 – 6 s) prevented the mice from turning based on temporal expectation of ICMS (**Figure 4B**). We ensured that turning was not random but was an ICMS stimulus-guided response by training the mice to suppress impulsive turns in the pre-stimulation period (**Figure 4B inset**). Representative psychometric curve (**Figure 4C**) shows that motivation was high throughout the task (100% responses to supra-threshold stimuli occurring at random trials), and impulsive and random turning was rare (close to 0% at sub-threshold stimuli). To efficiently and accurately measure behavioral detectability across multiple sites, we used an adaptive staircase method(Levitt, 1971), in which the amplitude of ICMS was raised or lowered based on the animal’s performance to estimate the threshold(Koivuniemi and Otto, 2011; Levitt, 1971) (STAR Methods, **Figure 4D**). This high-throughput method allowed us to quantify the threshold of 10 – 17 contacts spanning the cortical depth individually in one session. We performed the behavioral test longitudinally for up to 226 days in n=5 mice.

**Figure 4:**
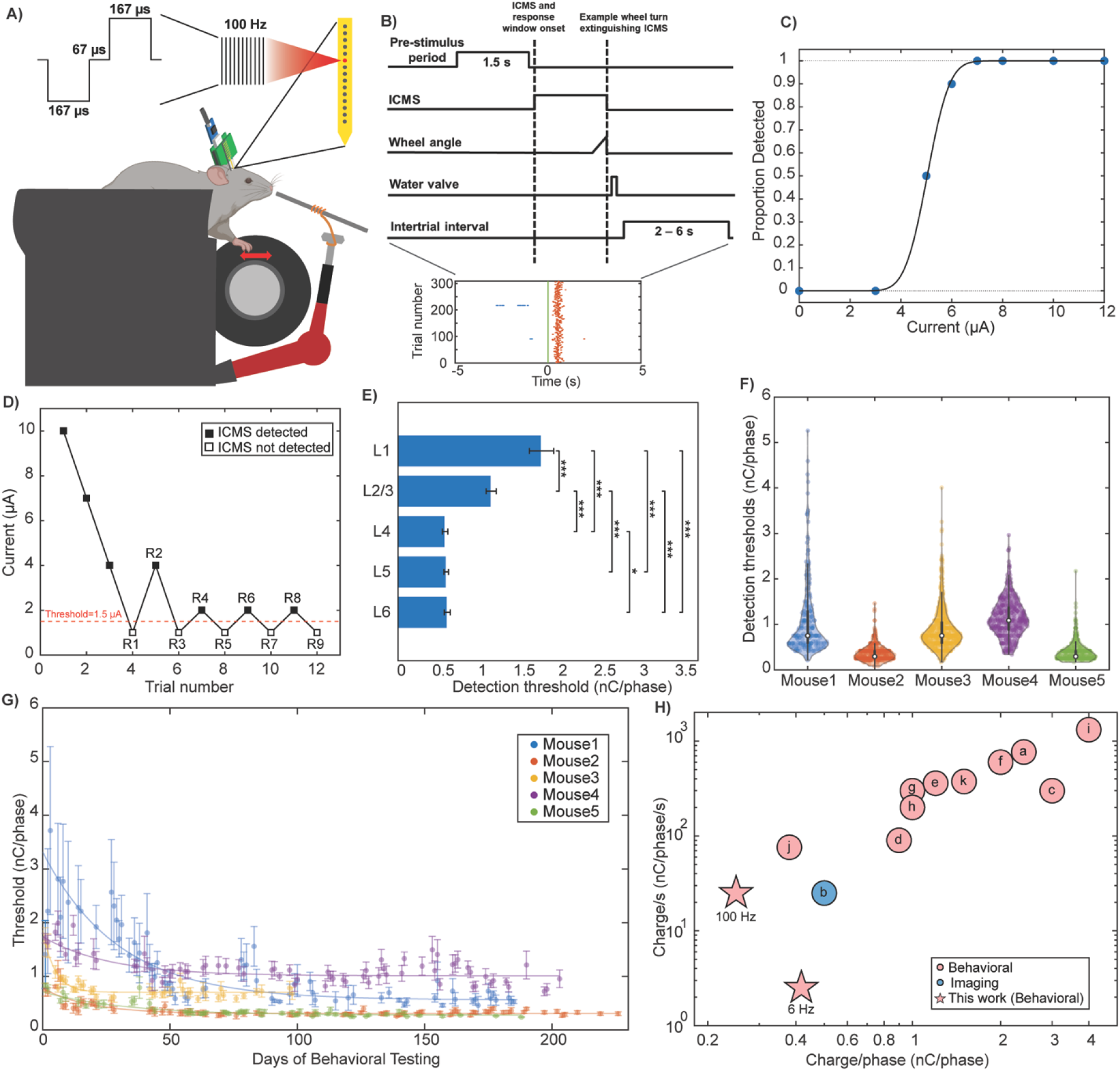
StimNETs elicit robust behavioral detection at low currents. **(A)** Sketch showing the wheel-turning task for ICMS behavioral detection. We used biphasic, cathode-leading pulses as depicted. The stimulation frequency was 100 Hz unless otherwise noted. **(B)** Diagram of trial structure used for the go/no-go task. ICMS was used both as the only cue and stimulus. Inset: response raster plot shows consistent, low latency response to suprathreshold stimulation (red dots) with very few impulsive turns in the pre-stimulation period (blue dots). t=0 marks the onset of ICMS. **(C)** Representative psychometric curve showing proportion of correct responses as a function of currents (n=1 session, 80 trials). **(D)** Representative threshold detection using adaptive staircase method. Threshold calculated as the average of last four reversals. **(E)** Detection thresholds at all cortical layers showing significant layer difference (Kruskal-Wallis test with Dunn’s post hoc correction, *** p<0.001, *p<0.05). n=5 animals, 64 stimulating contacts, and 362 sessions. Error bars denote 95% confidence intervals. **(F)** Violin plot showing averaged detection thresholds within cortical layers 4-6 for all mice across all sessions. n=5 animals, 64 stimulating contacts, and 362 sessions. **(G)** Detection thresholds of all contacts in L4 – L6 as a function of days showing lasting stability after initial decay. Solid lines are exponential fits. Error bars denote 67% confidence interval. n=5 animals, 64 stimulating contacts, and 319 sessions. **(H)** Literature comparison of ICMS behavioral detection (red) and neuronal activation (blue) threshold in rodents, non-human primates, and humans. Minimum reported or deduced values are plotted. A: Urdaneta et al. (2022)(Urdaneta et al., 2022), b: Zheng et al. (2022)(Zheng et al., 2022), c: Flesher et al. (2016)(Flesher et al., 2016), d: Hughes et al. (2021)(Hughes et al., 2021), e: Fernández et al. (2021)(Fernández et al., 2021), f: Callier et al. (2015)(Callier et al., 2015), g: Ferroni et al. (2017)(Ferroni et al., 2017), h: Ni et al. (2010)(Ni and Maunsell, 2010), i: Sombeck et al. (2020)(Sombeck and Miller, 2020), j: Schmidt et al. (1996)(Schmidt et al.), k: Rousche and Normann (1999)(Rousche and Normann, 1999).

We first concatenated all measurements from the entire experimental duration and examined the behavioral detectability as a function of cortical depth. We identified significant differences in thresholds across cortical layers (**Figure 4E,** Kruskal-Wallis test, χ2 = 707.84, p < 0.001, df = 4). Paired comparisons using Dunn’s post-hoc correction showed that shallow cortical layers L1, L2/3 had significantly higher detection thresholds than deeper cortical layers L4 – L6 (L1 vs. L4: p < 0.001; L1 vs L5: p < 0.001; L1 vs L6: p < 0.001; L2/3 vs. L4: p < 0.001; L2/3 vs L5: p < 0.001; L2/3 vs L6: p < 0.001) and L4 had the lowest threshold. The layer difference in detection thresholds was similar to other ICMS behavioral studies using rigid laminar probes(Urdaneta et al., 2021). Critically, the behavioral detection threshold by StimNET was low. **Figure 4F** shows the thresholds identified using all contacts in L4 – L6 from all measurement sessions (64 contacts, 362 sessions in total). The mean thresholds of all five animals were 1.12, 0.35, 0.88, 1.14 and 0.37 nC/phase, respectively, 3 out of which were lower than 1 nC/phase. The lowest single measurement thresholds of each animal were 0.21, 0.08, 0.17, 0.33, and 0.17 nC/phase, all of which were much smaller than 0.5 nC/phase. Particularly, the lowest measured value across subjects and sessions was 0.08 nC/phase (Mouse 2), which was at the limit of our stimulator’s resolution of 1 μA. The last four reversals of the staircase method occurred between 0 and 1 μA, resulting in a current threshold of 0.5 μA (**Supplementary Figure S1**).

We then scrutinized the time dependence of behavioral detectability of ICMS using StimNET contacts in deeper cortical layer of L4 – L6. Because contacts in L4, L5, and L6 provided relatively low detection thresholds, we analyzed all the data from L4 – L6 together without distinguishing the fine depth difference. In all animals, the detection threshold, averaged among all stimulation sites in L4 – L6, had an initial decay that can be described empirically as an exponential curve in the first 20 – 70 days. Then the threshold remained stable with little variation for a long-lasting period until we decided to terminate the experiment (**Figure 4G**). Going into finer details we also examined the stimulation currents in the last four reversals of the staircase method that led to the quantification of the threshold (**Supplementary Figure S2**). The currents in the last four reversals were stable within 1 μA (which is our stimulation current resolution) for n=78 sessions that spanned 226 days with a total of 1.9 million pulses output from this single contact.

The stable phase had much lower charge injection threshold than the initial phase during which the detection threshold decreased (**Figure 4G**). Notably, in two animals, a low current of 1.5 μA (0.25 nC/phase) was sufficient to maintain robust behavioral detection for 184 and 153 days, respectively, in the stable phase (**Supplementary Figure S3**). This provides the lowest threshold of chronic ICMS studies in either behavior detection or neuronal activation to the best of our knowledge. Furthermore, we explored if we could further reduce the overall charge injection by changing the stimulation frequency. We mapped the frequency-threshold dependence (**Supplementary Figure S4**) and found that by using a low frequency of 6 Hz, we reduced the charge injection per second by an order of magnitude at mildly elevated current threshold (**Figure 4H**). These results demonstrated that StimNET elicited robust, long-lasting, chronically stable behavioral detections at markedly low charge injections.

### StimNETs maintain tight tissue-electrode integration and normal function after chronic ICMS

We investigated the nature of the device-tissue interface by a combination of in vivo imaging and post-mortem immunohistochemistry. Representative examples of in vivo 2P imaging acquired two months after implantation showed dense, healthy vascular networks surrounding and in close contact with the implanted StimNET. Populations of neurons co-resided within microns of StimNET and the stimulating contacts with no signs of neuronal degeneration (**Figure 5A**). These observations are in qualitative agreement with the tight tissue-NET integration we reported previously without stimulation(He et al., 2022; Luan et al., 2017). To quantify the tissue response to the chronic implantation and stimulation of StimNET, we performed immunohistochemistry evaluations of the tissue surrounding StimNETs and compared between stimulating and passive (no stimulation) sites (**Figure 5B**). Fluorescence intensity of NeuN showed no changes with distance from StimNET, indicating the same neuronal density in the close vicinity of StimNET as far away. Fluorescence intensity of Iba-1 and GFAP had mild elevation within 50 μm of StimNET, but there was no encapsulation of microglia or astrocytes (**Figure 5B, C**). Critically, there were no differences between the stimulating and passive sites in any of these markers. These results suggest that StimNETs support the same stable, tightly integrated interface with brain tissue as the recording NETs, and that the simulating currents used in the study were within the safety limit and did not induce tissue damage.

**Figure 5:**
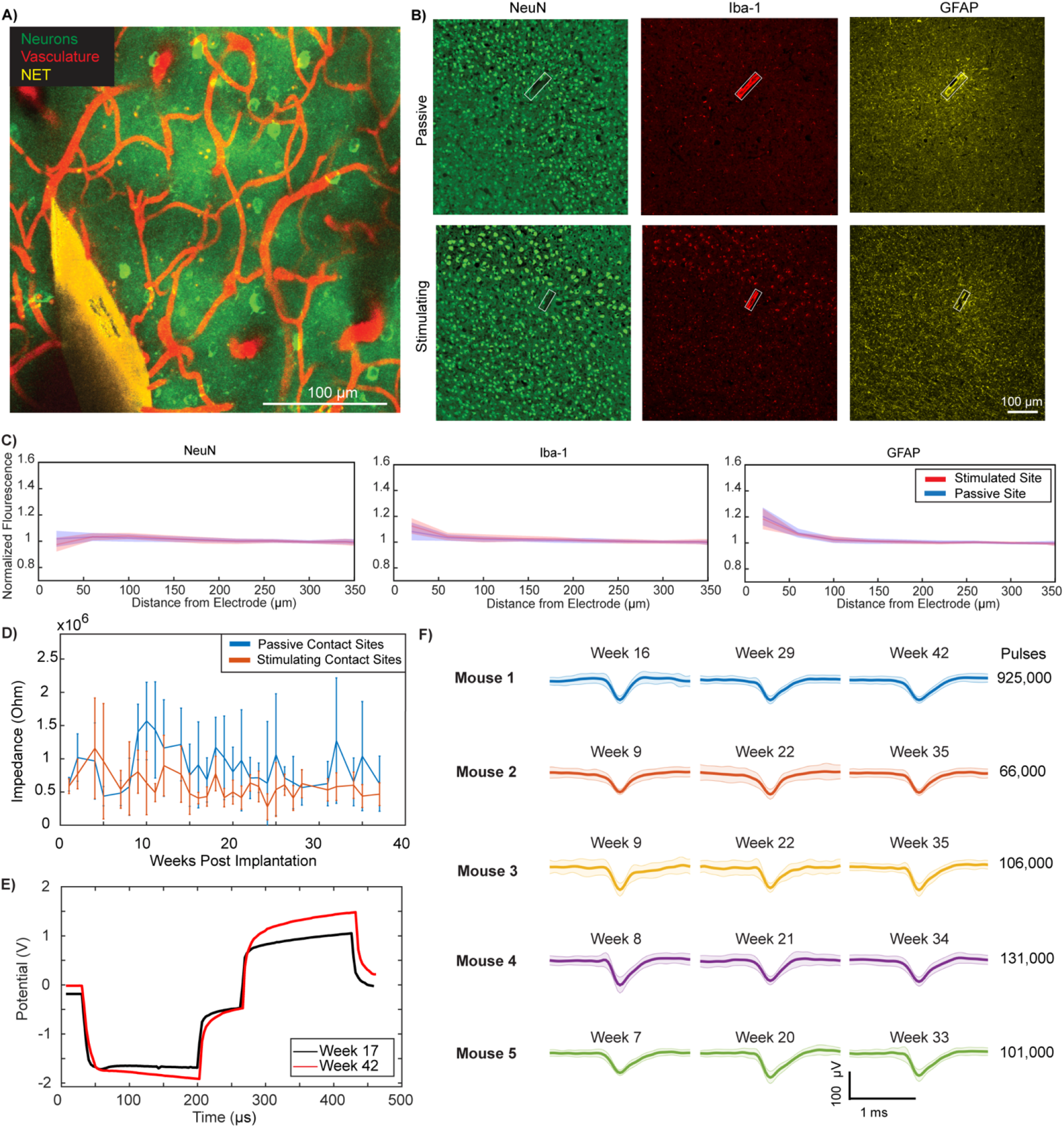
StimNETs maintain tight tissue-electrode integration and normal function after chronic ICMS. **(A)** Representative *In vivo* 2P MIP in a Gcamp6s mouse showing active neurons (green) and dense vasculature (red) around the StimNET. **(B)** Representative immunostaining for NeuN, Iba-1, and GFAP showing normal neuronal density and little glial scarring or aggregation around StimNETs both at the passive and stimulation sites. Green: NeuN; Red: Iba-1; Yellow: GFAP. Box encloses StimNET. **(C)** Fluorescence intensity as a functional of distance from StimNET of NeuN, Iba-1, and GFAP shows minimal disruption to local neuronal and glial cells from StimNET and no significant difference between stimulating and passive contact sites. Fluorescence intensity normalized to that from regions > 300 μm from StimNETs. (n=5 animals). **(D)** Chronic impedance at 1 kHz showed no significant changes over time for stimulating contacts (n=5 animals). Error bars denote 95th percentile confidence interval. **(E)** Voltage transients at Week 16 (Day 114) and Week 42 (Day 293) post implantation showing stability in charge injection after outputting 925,000 pulses *in vivo*. **(F)** Stimulation contacts recorded similar spike waveforms at similar SNRs months after stimulation. Stimulation pulse numbers of the contact site are noted.

Next, we investigated the functional integrity of StimNETs. The longest performing animal (Mouse 1 in the behavioral test) continued to behaviorally detect low amplitude stimulation after 308 days of implantation until its backend connector failed, underscoring the longevity of StimNETs *in vivo*. We analyzed the impedance of StimNETs as a function of days post implantation in the animals that underwent behavioral tests. Stimulation pulse number per animal was 4.7 million, 3.3 million, 2.4 million, 3.9 million, and 1.1 million (Mouse 1 – 5, respectively) with individual contact site pulse counts ranging from 12,000 to 1.9 million pulses. Impedance at 1 kHz showed stability over a chronic period of 37 weeks (260 days) and longer (to 44 weeks post-implantation with Mouse1) (**Figure 5D**). Furthermore, we repeatedly tested the change injection capability over the chronic periods of experiments. **Figure 5E** shows voltage transients in response to 8 μA biphasic pulses from a representative contact at Week 16 (Day 114) and Week 42 (Day 293) post-implantation (Week 0/Day 0 and Week 26/Day 179 after the first behavioral test) that had output over 925,000 pulses. There were no significant changes in the waveform shape or amplitude, further highlighting the chronic device stability of StimNETs for large amount of charge injections *in vivo*. Finally, we examined and verified that individual contacts on StimNETs recorded action potentials throughout the chronic stimulation periods. The waveform and the signal to noise ratio of extracellular spikes recorded from identical contacts remained the same following hundreds of thousands of in vivo stimulation pulses and after a chronic period of more than 8 months as they did within the first few weeks of implantation (**Figure 5F**).

## Discussion

Emerging neural electrode technology focusing on flexibility and miniaturization has made high-density, long-lasting, tissue-compatible neural recordings possible(Chung et al., 2019; Luan et al., 2017; Wei et al., 2018; Zhao et al., 2022). While it has been conceived that the same form factors that improved recording efficacy could also benefit intracortical microstimulation(Pancrazio et al., 2017), it remained challenging to make these small and flexible electrodes structurally and functionally robust for long-term stimulation. In this study, we engineered currently the thinnest, most flexible penetrating microelectrode array StimNETs for robust ICMS. Each microcontact on these devices stimulated up to 1.9 million pulses in vivo during 8+ months of intracortical implantation without signs of biotic or abiotic failures. The number of stimulation pulses StimNET output in vivo in this study was on par with a recent human study of ICMS that had a longer period of implantation^36^. These results suggested that StimNETs, with marked ultraflexibility and 1 μm total thickness, support long-term, clinically relevant applications of ICMS.

We leveraged the integrated application of in vivo imaging, behavioral, and histological techniques to decipher the spatial extent of neuronal activation, quantify the longitudinal behavioral detectability, and comprehensively characterize the tissue-electrode interface. Our application of multiple modalities stands out from previous studies where most often a single modality was employed and allows us to elucidate holistically the neuromodulation effects of StimNET-induced ICMS. We showed an intact and stable tissue-device interface of StimNET, low thresholds of 1 – 2 μA both by Ca^2+^ imaging and by behavioral detection, longitudinal stability in the neuronal activation and high level of spatial selectivity in time frames of a few months, and robust and stable behavioral detectability chronically up to 226 days. The neuronal activation threshold and the behavioral detection threshold are both among the lowest reported so far. In the stable phase of behavioral testing, the inter-day variations of median detectability in two animals (mouse 2 and 5) are at or smaller than 1 μA, which is the current resolution of the simulator used in the experiment, demonstrating superior longitudinal stability of StimNET. These results support our hypothesis that devices offering intimate tissue-electrode interface improve the focality, efficacy, and stability of ICMS and result in high-resolution, long-lasting, chronically stable neuromodulation. Critically, because StimNET elicited behavioral detections at very low currents of 1.5 μA, our approach provides an alternative path from current approaches of building more robust stimulators for large current stimulations and pursue a distinct regime of low current stimulation by tissue-integrated electrodes at little risk of interface deterioration or device abiotic failures from excessive charge injections.

Two-photon imaging has emerged as a powerful tool to decipher the neuronal response to ICMS at the single cell resolution(Histed et al., 2009; Michelson et al., 2019; Zheng et al., 2022). Most previous studies were performed acute (immediately after surgical procedures) and under anesthesia, which alters brain-wide neural activity including network effects (Friedberg et al., 1999). Rigid electrodes must be implanted at a large slant angle to accommodate the imaging objective, which may exacerbate the lasting neuronal process atrophy not only in the cortical depth dimension but also laterally along the cortical surface(Welle et al., 2020). Ultraflexible StimNETs can conveniently deflect for a 90 degree turn under the cranial window, allowing for steep implantation angles that are decoupled from the orientation of the carrier chip. We investigated neural responses to ICMS two weeks or more after the implantation, at which time the surgical trauma had subsided(He et al., 2022). Importantly, we performed all experiments in awake mice and developed a unique image processing workflow to quantify ICMS-evoked activity from the background of spontaneous activity at high throughput. These features allowed us to eliminate previous confounding factors and elucidate the spatial pattern of neuronal activation in the animal’s natural awake state at single-cell resolution, spanning sizable volumes, across various currents and over chronic periods. Our results at currents ≥ 7 μA showed similar, distributed activation of neurons as previous studies at same and larger current levels, which are consistent with neuronal activation of ICMS through the passage of axons^16^. Notably, at very low currents (2 μA) that did not elicit responses in most previous studies, we identified a more focal activation than larger, more commonly tested currents, which highlights the importance of lowering activation threshold to improving the spatial resolution of ICMS.

In one of the animals (Mouse 3) during longitudinal behavioral testing, a large current, estimated to be 40 μA, was accidently delivered for a few seconds. This incident immediately drove up the detection threshold that had been stable for 99 days at 4.45 ± 0.33 μA to 13.73 μA (**Supplementary Figure S5**). The detection threshold then subsequently decreased over a period of 116 days and finally settled at 4.87 ± 2.14 μA which was similar to the values prior to the incident. These results are consistent with our hypothesis that tight tissue-electrode integration is responsible for the low threshold we obtained. The change of detection threshold in this case can be explained as the large current we accidently delivered damaged the local tissue, which increased the average distance from excitable neurons to the stimulation site, so that higher currents were required to elicit the same behavioral response. The tissue healed over time, which lowered the average distance from neurons to StimNET and the detection threshold with it. These results restate the importance of healthy tissue-electrode interface for maintaining sensations at low stimulation currents without risking tissue safety.

The contact size used in this study is 24 μm in diameter, similar to what we developed previously for passive recording(Luan et al., 2017; Zhao et al., 2022) and warrants detection of single-unit activity as we demonstrated. While this study used single-shank, 32-channel devices, StimNETs are scalable owing to the wafer-scale microfabrication including sputter deposition of IrO_x_ and high-throughput implantation we have developed(Zhao et al., 2019; Zhao et al., 2022). Furthermore, StimNETs have the same miniaturized form factors, particularly the thickness, as their recording counterparts, which permits implantation of many these devices at a high volumetric density(Zhao et al., 2022). Taken together, StimNETs provide a scalable, long-lasting, chronically stable, bi-directional interface with neurons at high spatial resolutions.

## Acknowledgements

We thank Dr. K. Otto for helpful discussions; H. Rathore and Y. Jin for assistance with 2P imaging; Dr. S. Villapol for assistance with histological preparation; Dr. A. Budi Utama for assistance with histological imaging; the Animal Resource Facility at Rice University for animal care and housing, and Rice nanofabrication facility for the support on the microfabrication of StimNET. This work was funded by the National Institute of Neurological Disorders and Stroke, under R01NS109361 (to L.L.), and U01 NS115588 (to C.X.).

## Author contributions

Conceptualization, L.L. and C.X.; Methodology, R.L., R.K., and J.M.; Formal Analysis, R.L., R.K.; Investigation, R.L., R.K., and J.M.; Writing – Original Draft, R.L., R.K., and L.L.; Writing – Review & Editing, R.L., R.K., L.L., C.X., and N.T.; Funding Acquisition, L.L. and C.X.; Resources, R.L., P.Z., J.M., Y.S., A.K., E.A., B.N., R.Y., F.H.; Visualization, R.L. and R.K.; Supervision, L.L. and C.X.; The two co-first authors R.L. and R.K. have contributed equally to this work and are allowed to list themselves as first authors in their CVs.

## Declaration of interests

C.X. and L.L. and are co-inventors on a patent filed by The University of Texas (WO2019051163A1, 14 March 2019) on the ultraflexible neural electrode technology described in this study. L.L. and C.X. hold equity ownership in Neuralthread Inc., an entity that is licensing this technology. All other authors declare no competing interests.

## Inclusion and diversity

We worked to ensure sex balance in the selection of non-human subjects. One or more of the authors of this paper self-identifies as an underrepresented ethnic minority in their field of research or within their geographical location. One or more of the authors of this paper self-identifies as a gender minority in their field of research. One or more of the authors of this paper received support from a program designed to increase minority representation in their field of research. While citing references scientifically relevant for this work, we also actively worked to promote gender balance in our reference list.

## STAR Methods

### Resource Availability

#### Lead contact

Further information and requests for data and code should be directed to and will be fulfilled by the lead contact, Lan Luan.

#### Materials availability

This study did not generate new animal lines or unique reagents.

#### Data and code availability

All data reported in this paper will be shared by the lead contact upon request. All original code may be obtained from the lead contact upon request. Any additional information required to reanalyze the data reported in this paper is available from the lead contact upon request.

### Experimental model and subject details

#### Animals

A total of 14 mice at least 6 weeks of age or older, n=9 C57BL/6J-Tg(Thy1-GCaMP6s)GP4.3Dkim/J for 2P imaging experiments (4 male, 5 female) and n=5 (3 mice of C57BL/6J (1 male, 2 female) and 2 of GCaMP6s (1 male, 1 female)) for behavioral experiments were bred on-site from breeding pairs acquired from Jackson Laboratories (Bar Harbor, ME) and used in the experiments. Mice were single housed following implantation of StimNETs and housed within the Animal Resource facility at Rice University. 3 out of the imaging mice were excluded due to early occlusion of the cranial window and of breakage of backend connector. All surgical and experimental procedures in this study were in compliance with the National Institutes of Health Guidelines for the Care and Use of Laboratory Animals and were approved by the Rice University Institutional Animal Care and Use Committee.

### Method details

#### StimNET fabrication

The 32-channel, single-thread StimNETs were fabricated by conventional photolithography and metallization on fused silica wafers using a multi-layer structure. The microfabrication procedure had the following steps. i) A nickel metal release layer was patterned by depositing 3nm Ti and 60nm Ni under the flexible section of the device. ii) A bottom insulation layer was created by spin coating a diluted polyimide polymer (PI2574, HD Microchemicals) to reach ~500 nm thickness and baked in a vacuum oven according to the manufacturer’s recommended thermal profile. iii) An interconnect layer was defined by photolithography and metallization of a 3nm Cr, 100nm Au, and 3nm Cr metal stack by electron (e)-beam evaporation (Sharon Vacuum Co., Brockton, MA). Additional layers of 3 nm Cr, 160 nm Ni, and 80 nm Au were deposited on the solder pads to increase the reliability of solder reflow and reduce alloying of solder and gold. iv) The top insulating layer was created in the same method as the bottom layer. v) The thread outline, via to the electrodes, and solder pads were defined by RIE etching (Oxford Instrument) using O_2_/CF_4_ gas mixture in the 9:1 ratio. vi) Microcontacts for recording and stimulation were defined by photolithography and sputter coating of 10nm Ti, 100nm Pt, 10nm Ti, and 300nm IrO_x_ stack (AJA ATC Orion Sputter System). vii) A capping layer of 300 nm PI is defined and etched as described in previous steps. viii) Low-temperature solder balls were placed on solder pads to form a ball grid array using a solder jetting tool (PacTech), and the wafer was diced into individual devices. The StimNETs were then individually bonded to a custom printed circuit board (PCB) to interface with recording/stimulation electronics. Then the flexible section of StimNETs was released from the substrate by etching of the Ni layer, and the glass substrate was cleaved to the desired length. Lastly, the flexible implantable potion of the StimNET was affixed to a 50-μm diameter sharpened tungsten wire via Polyethylene glycol (PEG), which served as a temporary adhesive securing the probe for implantation as previously described in detail(Yin et al., 2022).

#### Simulation of electric stimulations

Finite element (FE) model simulations of the electrical stimulation produced by StimNET electrodes were conducted in COMSOL Multiphysics 5.6 (COMSOL, Inc., Burlington, MA). StimNETs were modeled in COMSOL with dimensions matching those utilized in this study, implanted within the center of a uniform block (1.5 × 1.5 × 3 mm) of neural tissue. To evaluate the impact of a glial scar on stimulation efficacy, an encapsulating volume representing the glial scar of 0, 20 and 40 μm was used. The FE models contained between 14,414,413 and 14,564,731 elements depending on glial scar thickness, and used following electrical properties: PI substrate, conductivity of 1e-12 S/m and permittivity of 11.7; metal contact sites, conductivity 9.43e6 S/m and permittivity of 2.7604; neural tissue, conductivity 0.2 S/m and permittivity of 88.9; glial scar, conductivity 0.166 S/m and permittivity of 88.9.

To quantify the population of neurons activated by monopolar stimulation, the stimulation microcontact was modeled as the current source and the outer boundaries of the model as the ground. The current injected through the source was varied, and the volume of activated tissue was measured to be the neuronal tissue, excluding the glial scar, that reached or exceeded a charge density threshold of 1292 μA/mm^2^ following values from reference(Tehovnik et al., 2006). A density of neurons of 110,000 neurons/mm was used(Keller et al., 2018) to quantify the number of neurons activated by this stimulation. The effect of glial scar thickness was examined by running the simulations with no glial scar element or including either a 20 or 40 μm thick scar.

To quantify the spatial selectivity of neuronal tissue activation, two neighboring contact sites with an inter-site distance of 60 μm (center-to-center) at varied stimulation currents and with glial scars of 0, 20, or 40 μm thick were modeled using the identical conditions as above. The volume of neuronal tissue activated from each contact independently was referred to as the ‘single electrode stimulated volume’, and the volume of activated neuronal tissue activated by both contact sites was referred as the overlapping region.

#### In vitro characterization of StimNET

The charge injection and storage capacity of StimNETs was evaluated by cyclic voltammetry (CV) and voltage transient measurements in saline using Gamry Reference 600+ (Gamry Instruments, Warminister, PA). Measurements were made in a three-electrode setup using a large-area platinum counter electrode and Ag/AgCl (3M NaCl) reference electrode (BASi Research Products, West Lafayette, IN). Voltage transients were measured in response to biphasic pulses with 100 μs pulse width, and 67 μs interphase interval of various amplitudes. CV measurements used a sweep rate of 100 mV/s and were swept between 0.8V and −0.6V.

#### Surgical procedure

All animals received co-implantation of a cranial optical window and StimNET in one surgery as described previously (Yin et al.). Briefly, animals were anesthetized with isoflurane (3% for induction and 1%-2% for maintenance) and administered extended-release Buprenorphine (Ethiqa TM) and Dexamethasone (2 mg/kg, SC) for analgesia and to reduce surgery-induced inflammation, respectively. The surgical site was infiltrated with lidocaine (7 mg/kg 0.05%) subcutaneously prior to shaving and disinfected with 3 × iodine and alcohol wash before the initial incision into the skin of the head. The skull was exposed between bregma and lambda skull sutures, followed by the removal of the fascia and scoring of the skull crosshatch pattern to prepare the skull. A circular craniotomy of dimensions 3mm in diameter over the somatosensory cortex was drilled in the skull for the StimNET implantation, and a burr hole was drilled in the contralateral hemisphere to accommodate a Type 316 stainless steel grounding wire. Following the opening of the craniotomy, a 32-contact StimNET affixed to a 75 μm tungsten wire via the bio-dissolvable adhesive PEG was implanted through the dura to the somatosensory cortex by stereotaxic targeting at approximately 2 mm ML and −1.5 mm AP at an insertion angle of 30 degrees off vertical, though variations in exact position were made to accommodate surface vasculature and ensure a clear region in the vicinity of the probe implantation site to permit imaging. Following the implantation of the StimNET, the PEG affixing the StimNET to the shuttle wire was allowed to dissolve, and the wire was removed. A sterile glass coverslip window (#1, manufacturer) was secured over the craniotomy using cyanoacrylate adhesive and Metabond dental cement (Parkell, NY) with regions not directly covered by glass filled with Kwiksil (World Precision Instruments). Additional dental cement was applied to adhere a headbar for head fixation to the skull cap and seal the cranial window to the skull. Animals were provided at least three days of recovery post-surgery and an additional three days of familiarization to head restraint before the beginning of any experimental tasks.

#### Two-photon imaging

Two-photon (2P) imaging was performed using a laser scanning microscope (Ultima 2p plus Bruker, MA) equipped with a 16’ water immersion objective (numerical aperture of 0.8, Nikon, NY) and an ultrafast laser tuned to 920 nm for fluorescence Ca^2+^ excitations (InSight X3, Spectra-Physics). After initial habituation, z-stack 2P imaging was performed, for which mice were awake and head-restrained on a home-constructed low-friction rodent-driven belt treadmill following the design of HHMI Janelia (https://www.janelia.org/open-science/low-friction-rodent-driven-belt-treadmill). Each imaging session lasted up to 3 hours and contained multiple replicants. Each replicant contained alternating stimulation and baseline (no stimulation) trials from randomized stimulation sites and currents. 512 × 512 images of a field of view up to 1 mm × 1 mm were acquired at 30 fps using galvo-resonant scanners. The duration of a typical z-stack, referred to as an imaging trial, was about 30 s for a depth of 400 μm at the z-spacing of 2 μm. An inter-trial period of 2 to 5 seconds was implemented for data saving. Electrical stimulation was delivered via a custom Pico32+Stim front end with a Grapevine neural interface processor (Ripple Neuro, Salt Lake City, UT). During stimulation trials, 50 Hz electrical stimulation pulse trains of biphasic, charge-balanced cathode-leading square pulses at 167 μs per phase and 67 μs inter-phase interval were provided. The current amplitudes were 2, 5, 7, and 10 μA, resulting in a maximum charge injection of 1.67 nC/ph per phase and a maximum charge density of 369.5 μC/cm^2^. According to the Shannon criteria, the largest stimulation current gave a K=0.48, smaller than the threshold of 1.85 for tissue compatible/safe neural stimulation. Customized MATLAB (MathWorks, MA) scripts were developed to randomize stimulation parameters and control data acquisition. Stimulation and 2P imaging were synchronized via TTL signals generated by a PulsePal (Sanworks, NY) unit.

#### Behavioral training

After a post-surgical recovery period of 7 days, the animals undergoing behavioral testing were put on water restriction Monday through Friday and on ad libitum water during the weekends and holidays. The animals were monitored every weekday to ensure their weights were above 85% of baseline body weight. Every animal received a minimum of 1 mL per weekday. If the animal did not receive all its daily allotment of water during the behavioral task, the remainder of its daily allotment was given after an hour had passed following the end of the behavioral session. Behavioral testing was performed using a standardized experimental rig from the International Brain Laboratory(International Brain et al., 2021) with the following customizations. To accommodate electronics and cabling of StimNET, custom-fabricated headbar holders were used. The wheel for decision making was oriented 90 degrees such that the wheel could be spun forwards and backward rather than left and right, allowing for a more natural movement for a go/no-go task. Electrical stimulation was delivered via a Pico32+Stim front end customized for small current output with a Grapevine Neural Interface Processor (Ripple Neuro, Salt Lake City, UT) onto StimNET sites that had impedance < 1 MΩ at 1 kHz. All microstimulation was performed using cathode-leading pulses with a pulse width of 167 μs and 67 μs interphase interval. The frequency of stimulation was kept at 100 Hz. The stimulation and recording were controlled via Xippmex MATLAB application programming interface (The MathWorks Inc., Natick, MA) on a computer separate from the behavioral task controlling computer.

The behavior training had two stages after acclimation to handling and head fixation. At Stage 0, the animal freely turned the wheel without stimuli and obtained a water reward (10% sucrose solution) for wheel-turning behavior every trial passing an angular threshold. The initial angular threshold started at 20 degrees and increased with sessions and response rate to a final value of 30 degrees. The animal graduated stage 0 training once the response rate exceeded 95%. The purpose of this stage was to shape goal-directed behavior by forming a response-outcome association between the wheel turn and sugar water. Stage 1 introduced single-site suprathreshold (15 μA) ICMS as the stimulus and the response-outcome association was made contingent upon the stimulus to form a stimulus-response-outcome chain. At this stage, the animal was rewarded by turning the wheel past the angular velocity threshold (i.e., a Go response) during a response period beginning after the stimulus. The response period had an initial duration of 10 s and was concurrent with a 10 s stimulus. A Go response during the response period resulted in extinguishing the stimulus and reward delivery. If the animal did not respond within the response period, there was no penalty. Each trial was followed by an intertrial interval (ITI). The ITI was randomized and drawn from a uniform distribution with an initial interval of 2 to 3 s. The trial began with a pre-stimulus period (PSP) of 0.5 s to discourage premature responses. Responses during the PSP were negatively reinforced by resetting PSP, which delayed the stimulus and thus delayed opportunity to receive reward. If the animal appeared to respond well to the suprathreshold stimulation with PSP violation rate less than 30% of trials, the response period was decreased incrementally (1-2 s) over multiple sessions to a final period of 1 s. Similarly, the ITI upper bound was incrementally increased (0.5-1 s) to 6 s such that the final ITI was drawn from a uniform distribution with an interval of 2 to 6 s. PSP was incrementally increased (0.2-0.3 s) to a final value of 1.5 s. If the animal did not respond to stimulation, the animal was given one more session stimulation before another site was chosen. After the animal was able to produce consistent low latency responses within 1 s with a low PSP violation rate (<10% of trials), the animal graduated training and advanced to detection threshold measurements. Animals typically needed 2 weeks of training to proceed to detection threshold measurements.

#### ICMS threshold detection

To measure the ICMS detection thresholds across multiple contact sites and animals efficiently, an adaptive staircase procedure was employed. This procedure was run for each of the viable stimulation sites in a randomized manner. For a given trial, if the animal responded to the stimulus, the current for the subsequent trial was decreased by a step of 1 μA. However, if there was no response, the current for the next trial was increased by a step of 1 μA. The staircase procedure terminated if it did not respond to the maximum current (25 μA) trials three consecutive times, the number of trials exceeded 25 trials, or after nine reversals where a reversal is defined as the transition from an increasing or decreasing trend to a subsequent decreasing or increasing trend, respectively. The threshold was the average of the last four reversals. The initial step size was 3 μA and changed to 1 μA after the third reversal.

#### In vivo electrophysiology

Voltage transients were measured with respect to a Type 316 stainless steel reference wire. Voltage transient measurements were performed weekly on behavioral animals using the chronopotentiometry function on the Gamry Reference 600+ (Gamry Instruments, Warminister, PA) to assess the functionality of the stimulating electrodes. The input current waveform was a cathode-leading biphasic charge-balanced pulse with a pulse width of 167 μs and 67 μs interphase interval with an amplitude of 8 μA. The maximum cathodically and anodically driven electrochemical potential excursions (E_MC_) were measured as the potential 20 μs after the end of the cathodic and anodic phases, respectively.

In vivo impedance measurements at 1 kHz were performed weekly on animals using an Intan RHS stim/recording controller and RHS 32-channel stim/recording headstage (Intan Technologies, Los Angeles, CA). Contact sites which report impedances over 3 Mohm are considered to have a broken backend connection and removed from subsequent impedance measures. Neural electrophysiological recording was performed intermittently on the animals under behavioral training using Intan or Ripple. For spike detection, raw data were filtered with a bandpass filter with lower and upper cutoff frequencies of 300 and 5000 Hz. Spike detection was performed using the MATLAB command “findpeaks” to find spikes. Since “findpeaks” looks for positive-valued peaks, the sign of the input signal was flipped. The threshold value was set to 60 μV. The minimum distance between peaks was set to 1.5 ms. Additionally, the “halfprom” minimum peak prominence, maximum peak width, and minimum peak width were set to 40 μV, 25 samples, and 7 samples, respectively. Dimensionality reduction was performed with PCA by keeping the top two variance-explaining components. K-means clustering was subsequently applied to the 2-dimensional data points with k=3. If the mean of each cluster resembled a neural spike via manual inspection, the largest amplitude cluster was kept and plotted. Clusters that failed manual inspection were discarded.

#### Histological tissue collection and analysis

For animals implanted for more than three months, brain tissue was collected and histologically processed to quantify the chronic immune response to implanted StimNETs. For perfusion and tissue collection, animals were first anesthetized with isoflurane (3%-5%) and perfused PBS transcardially at 80 mmHg through the circulatory system of the animal until outflow was clear, followed by a fixative solution of roughly 500 mL 4% paraformaldehyde; both fluids were chilled to 4 °C. Following perfusion, the head was removed from the body, burr holes were drilled throughout the skull to improve fluid flow, and the head was placed in a 4% paraformaldehyde solution for 48 hours at 4 °C for fixation. Afterwards, the heads were cryoprotected by immersion in a 10% sucrose solution for 72 hours and then frozen at −80 °C for at least 24 hours before extracting the brain from the skull. Care taken to ensure the implanted electrode was not mechanically disturbed during the skull extraction. The frozen brains were then cryosectioned at 20 μm thick via cryostat (CM1520 Leica Biosystems, IL), and slices were transferred to 48-well cell culture plates for fluorescence labeling.

Tissue slices were prepared for histology by first rinsing tissue three times in a 1 × PBS solution with 0.1% triton X-100 for 5 minutes each before blocking tissue in a 10% BSA solution for 1 hour with gentle agitation at room temperature. Slices were then incubated with primary antibodies for neurons with conjugated mouse anti-NeuN (1:100 dilution, MAB377X; Millipore), microglia with chicken anti-GFAP (1:5000 dilution, ab134436; Abcam), and astrocytes with conjugated rabbit anti-Iba1 (1:1550 dilution, 015-28011; Fujifilm) in a solution containing 1% BSA overnight at 4 °C. Slices were again rinsed three times in a 1 × PBS solution with 0.1% triton X-100 for 10 minutes each before being incubated in secondary antibody Goat Anti-Chicken Alexa Fluor^®^ 647(1:200 dilution, ab150171; Abcam) for 1 hour at 4 °C. Slices were washed three more times in a 1× PBS solution for 5 minute periods before being sealed with Vectashield plus antifade mounting medium (Vector Laboratories, CA) doped with DAPI. Slides were placed in a dark chamber at 4 °C for at least 24 hours before imaging.

Confocal imaging of brain slices was performed with a Nikon A1 confocal microscope (Nikon Instruments Inc., Melville, NY). Four fluorescent channels were imaged to quantify the tissue response of neurons (488 nm), microglia (641 nm), astrocytes (561 nm), and cellular nuclei (405 nm). To quantify fluorescence intensity as a function of distance from StimNET, the boundary of StimNET in each tissue section were manually outlined. Then a custom analysis script written in ImageJ(Schneider et al., 2012) and MATLAB (MathWorks, MA) defined contours every 40 μm from 0 – 360 μm from the StimNET boundary. For each fluorescence channel, the average fluorescence in areas between every two adjacent contours was calculated as the intensity at that distance away from StimNET and normalized against the average fluorescent intensity in areas of 280 – 360 μm away from StimNET.

### Quantification and statistical analysis

#### Identification of stimulation-induced Ca^2+^ neural activation

2P imaging data were processed using a custom-written program integrating MATLAB (MathWorks, MA) and ImageJ(Schindelin et al., 2012) to identify and isolate neurons activated by neural stimulation from the background (**Fig 2D**). The first goal is to identify regions of interest representing activated neurons in an imaging session. First, the voxel-by-voxel value of fluorescence intensity and standard deviation (STD) was calculated for all the baseline (no stimulation) trials during an imaging session, which provided a quantification of the spontaneous (passive) neural activity of the brain. Next, the stimulation-induced fluorescence increase was determined by subtracting baseline fluorescence from stimulation trials voxel by voxel. The baseline fluorescence was determined by averaging six baseline trials temporally proximal to the stimulation trial, the three prior and three after, to minimize the variability in individual baseline scans. The stimulation-induced activation was then determined as binarized voxels at a threshold of stimulation-induced fluorescence increase greater than three times the STD of the baseline fluorescence. These steps were performed for all stimulation trials, and the regions of stimulation-induced activations were summed across all trials to generate the voxel-by-voxel map containing all regions of activation that were weighted by the number of trials of activation. Finally, all regions of activation were segmented by the ImageJ plugin ‘3D iterative segmentation’(Ollion et al., 2013) to provide regions of interest (ROIs) that defined neurons activated throughout the entire imaging session across stimulation sites and currents.

The next goal is to identify neurons activated by a particular stimulation parameter. First, randomized stimulation scans were re-grouped by stimulation sites and currents. For each stimulation parameter, the same differential calculation as described above was repeated to obtain maps of stimulation-induced fluorescence increase, which was then masked by the segmented ROIs obtained previously and binarized by the same threshold as previously discussed to obtain stimulation-activated neuron ROIs. Then consistency of activation was checked across N trials under identical stimulation parameters, and ROIs consistently activated by more than 75% were marked as stimulation-activated neurons.

#### Quantification of neuronal activation via 2P imaging

To evaluate the neuronal response to ICMS, several metrics were quantified after the identification of activated neurons by 2P imaging, including the number of activated neurons, the consistency of activated neurons, the distance of neural activation, the density of activation, and the spatial specificity of stimulation. Calculating the number of activated neurons for a given stimulation current or population was accomplished by simply summing the total number of activated neuron ROIs detected. The consistency of neural activation across currents was calculated by tracking the activation pattern for individual neuron ROIs from a lower stimulation current to the next higher (e.g., 2 μA to 5 μA) and dividing the number of neurons activated at both currents by the total number of neurons active at the lower current level. The maximum three-dimensional neural activation distance from the stimulating contact site was considered the maximum Cartesian distance between the centroid of an activated neuron and the stimulating contact site. The activation density was calculated as the number of activated neurons detected within 100-μm thick spherical shell bins emanating radially from the stimulating stimulation divided by the volume of the shell within the imaging volume (**Supplementary Figure S1**). The spatial selectivity of activation was calculated by comparing the populations of neurons activated by two nearby sites within the same imaging session.

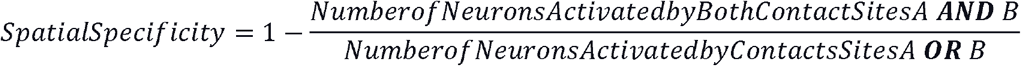

Spatial specificity was evaluated for contact sites spaced 60, 120, and 180 μm apart and across all stimulation current levels.

#### Statistical analysis

All statistical analyses were performed in MATLAB (MathWorks, MA). Kruskal-Wallis tests with Dunn’s post-hoc were used to compare the population of activated neurons by stimulation current, comparison of the distance of neural activation by stimulation current, the selectivity of stimulation at varied currents and distances, and the detection threshold of stimulation across cortical layers. Welch’s t-tests were employed for the pairwise comparisons at each binned distance for the histological evaluation of tissue neighboring stimulating versus passive contact sites.

## Notes

### Competing Interest Statement

L. L., C.X., and P.Z. hold equity ownership in Neuralthread Inc., an entity that is commercializing the ultraflexible electrode technology described herein. All other authors declare no competing interests.

