## Supplementary materials for "Low-threshold, high-resolution, chronically stable intracortical microstimulation by ultraflexible electrodes"

**Supplementary Figures:**

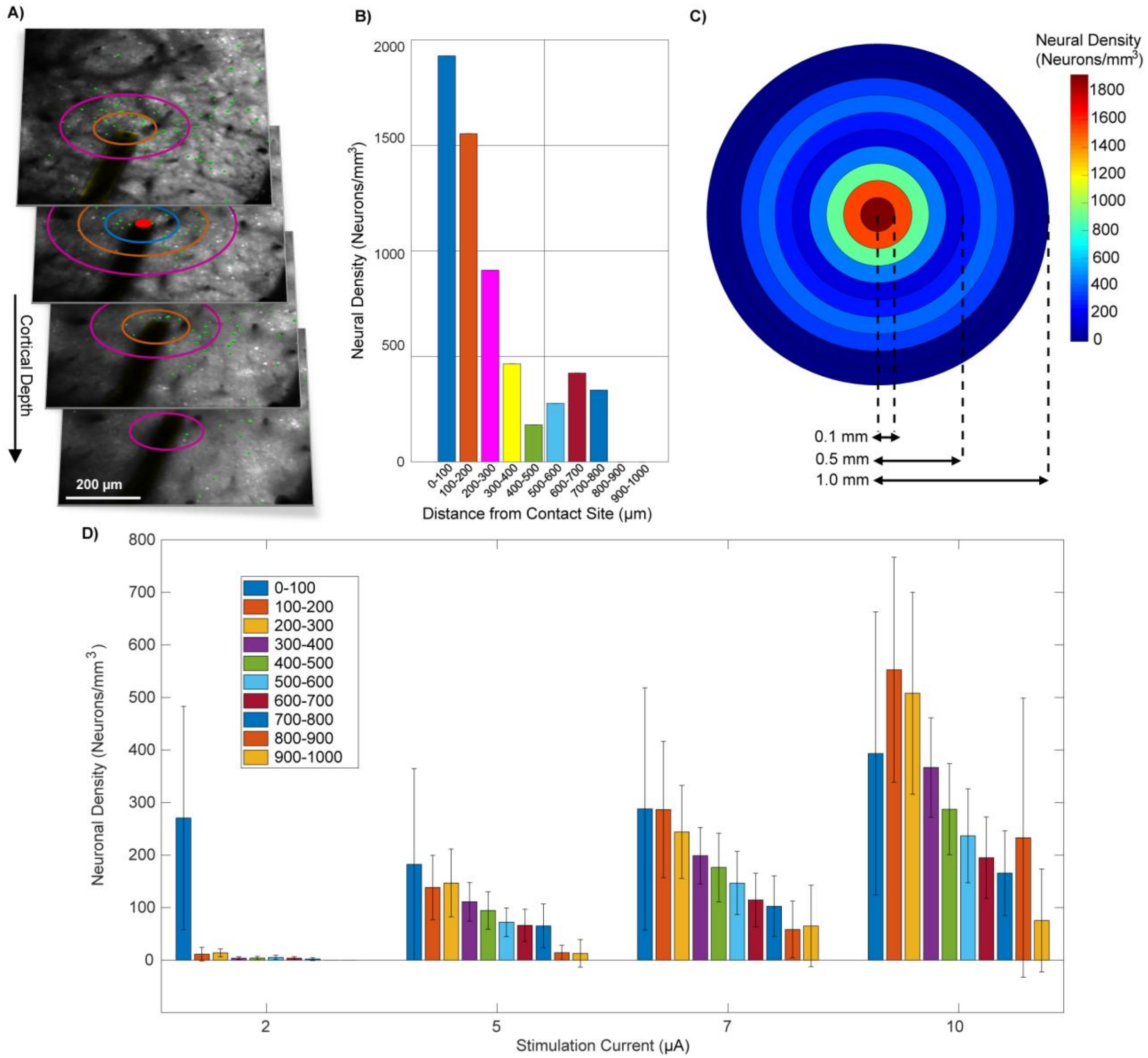


**Supplementary Figure S1: Quantification of volumetric neural activation density**. **(A)** Representative z-stack showing activated neurons (green) and their distance to the stimulation site (red dot). Color coded rings at each image denotes the Cartesian distance in 3D every 100 μm (purple; 300 μm, orange: 200 μm; blue: 100 μm). Neurons enclosed between two rings contribute to the activation density at the matching distance from the stimulation site as shown in (B). Note that separation along cortical depth z is taken into account in calculating the distance, so that the ring size of a specific radius from the stimulation site reduced as the imaging plane moved away from the stimulation plane. **(B)** Representative bar plot of density of activated neurons as a function of distance from the stimulating site. Same color code as in (A). In (A) and (B), stimulation depth is 100 μm; stimulation current is 10 μA. **(C)** Example ring plot generated from (A) and (B) describing the density of evoked neurons as a functional of distance from the stimulation sites via color coded heatmap. (**D)** Bar plots of density of activated neurons as a function of distance from the stimulating site for all four levels of currents, 2, 5, 7, and 10 μA. Data averaged 5 animals, 11 imaging sessions, and 21 stimulation sites (same as Figure 3C)

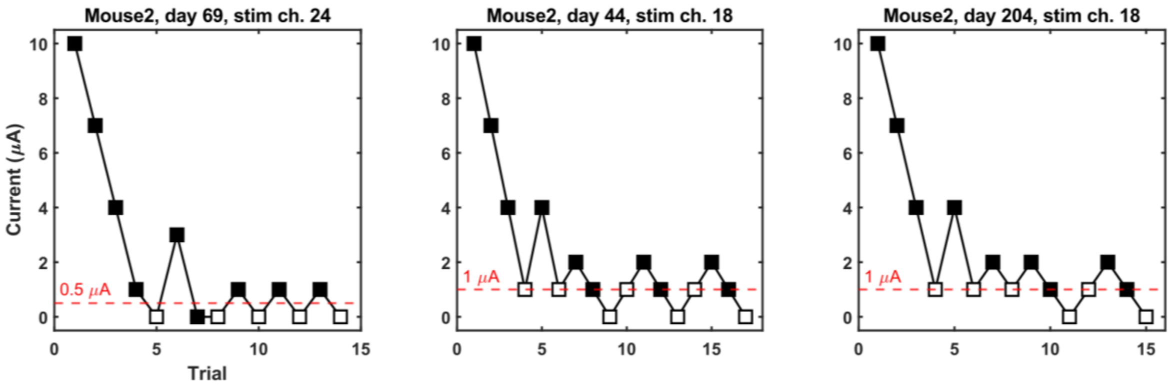


**Supplementary Figure S2: Examples of staircases with lowest detection thresholds.** Staircases showing lowest measured detection thresholds of 0.5, 1, 1 μA respectively. Black square indicates stimulus detected while white square indicates stimulus was not detected. The threshold (red dotted line) is the average of the last four reversals. Initial step size is 3 μA with step size of 1 μA after the third reversal.

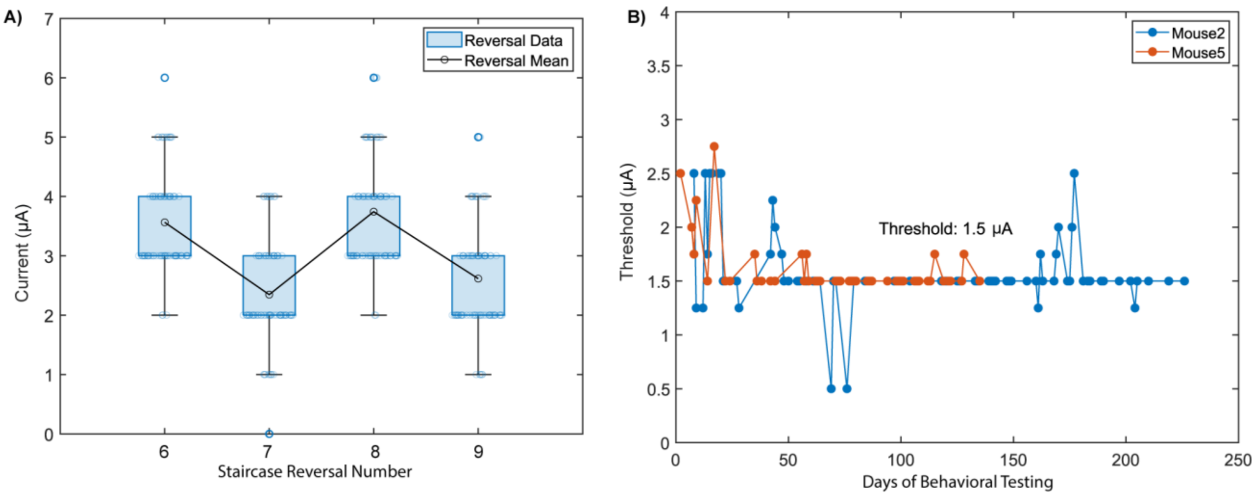


**Supplementary Figure S3: Longitudinal tracking of last four reversals and low detection thresholds demonstrates stability. (A)** Last four staircase reversals for all sessions (n=78) of a representative stimulation contact with 1.9 million stimulation pulses. **(B)** Additionally, the behavioral detection threshold of a single contact from two animals remained at a markedly low level of 1.5 μA for up to 230 days into behavioral testing. Experiments in Mouse 5 had to terminate earlier due to connection failures.

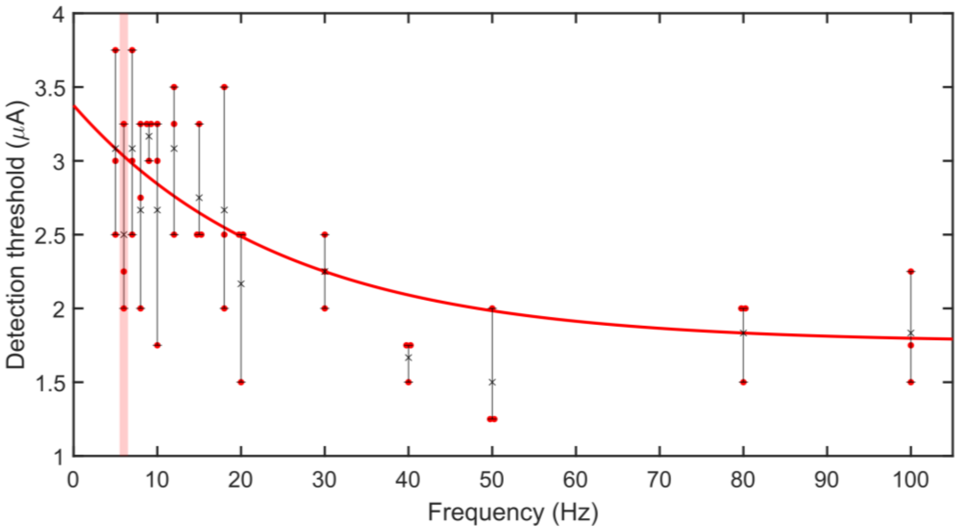


**Supplementary Figure S4: Minimizing charge per second detection threshold.** Detection threshold vs. stimulation frequency over 3 sessions. Error bars range from minimum to maximum of the three measured thresholds for a given frequency. The X marker represents the mean. A decaying exponential was fit to the mean thresholds. The red shaded bar indicates the frequency (6 Hz) that minimizes charge per second (nC/s) for this specific stimulation contact.

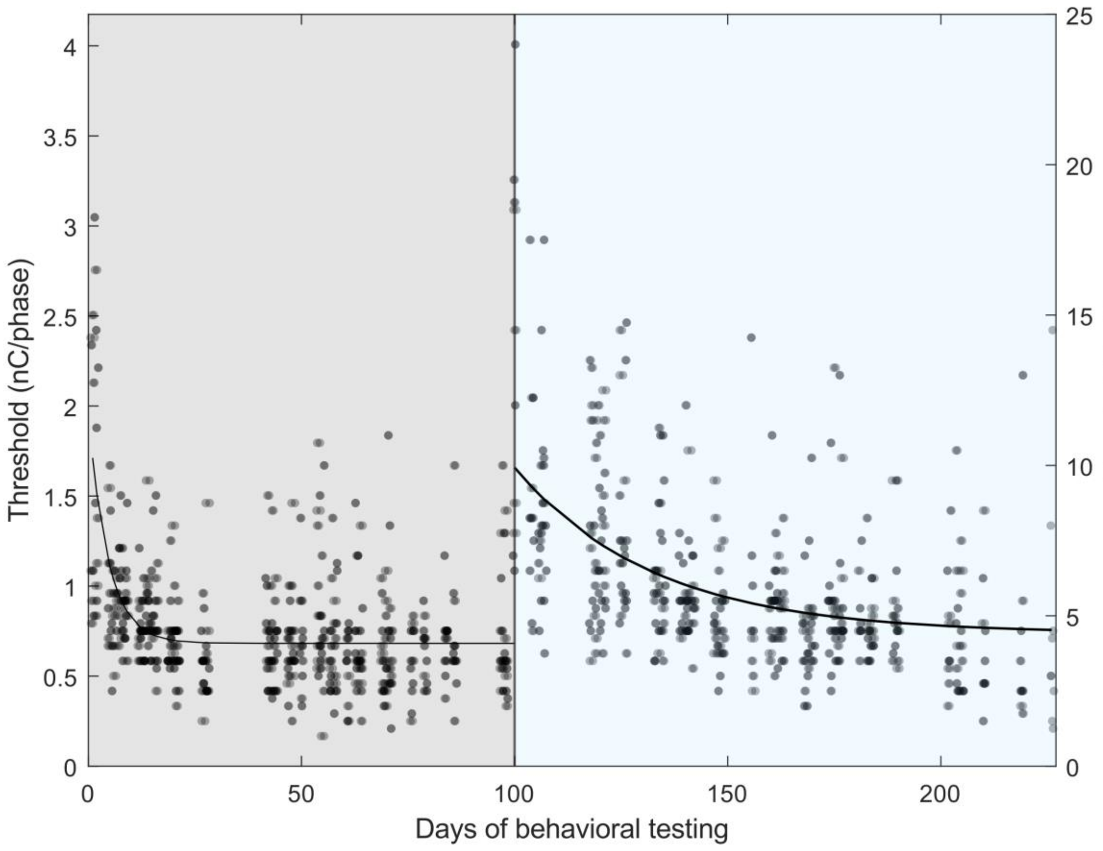


**Supplementary Figure S5: A single-time high stimulation direct current causes elevated thresholds that subsided with time.** Detection thresholds before the accidental 40 μA direct current stimulation (gray) and after (blue) for Mouse 3. Asymptotic thresholds for pre-shock and post-shock segments were calculated by grouping the last four sessions from each segment (n=12 contacts pre-shock, 9 contact post-shock) with no significant difference found between the two groups (Kruskal-Wallis test, $\chi^2 = 0.40$, $p = 0.4$, $df = 1$). Days are with respect to the start of threshold measurements. Each point represents the threshold for an individual site for a given session. A decaying exponential curve was fitted to the mean thresholds for both segments. A horizontal jitter amount of 0.5 was applied to minimize data point overlap.
